# Single-cell analysis reveals inter- and intratumour heterogeneity in metastatic breast cancer

**DOI:** 10.1101/2022.10.16.512415

**Authors:** Baptiste Hamelin, Milan Obradović, Atul Sethi, Michal Kloc, Simone Muenst, Christian Beisel, Katja Eschbach, Hubertus Kohler, Savas Soysal, Marcus Vetter, Walter P. Weber, Michael B. Stadler, Mohamed Bentires-Alj

## Abstract

Metastasis is the leading cause of cancer-related deaths of breast cancer patients. Some cancer cells in a tumour go through successive steps, referred to as the metastatic cascade, and give rise to metastases at a distant site. We know that the plasticity and heterogeneity of cancer cells play critical roles in metastasis but the precise underlying molecular mechanisms remain elusive. Here we aimed to identify molecular mechanisms of metastasis during colonization, one of the most important yet poorly understood steps of the cascade. We performed single-cell RNA-Seq (scRNA-Seq) on tumours and matched lung macrometastases of patient-derived xenografts of breast cancer. After correcting for confounding factors such as the cell cycle and the percentage of detected genes (PDG), we identified cells in three states in both tumours and metastases. Gene-set Enrichment Analysis revealed biological processes specific to proliferation and invasion in two states. Our findings suggest that these states are a balance between epithelial-to-mesenchymal (EMT) and mesenchymal-to-epithelial transitions (MET) traits that results in so-called partial EMT phenotypes. Analysis of the top Differentially Expressed Genes (DEGs) between these cell states revealed a common set of partial EMT Transcriptions Factors (TFs) controlling gene expression, including ZNF750, OVOL2, TP63, TFAP2C and HEY2. Our data suggest that the TFs related to EMT delineate different cell states in tumours and metastases. The results highlight the marked interpatient heterogeneity of breast cancer but identify common features of single cells from five models of metastatic breast cancer.

## Short Intro

Breast cancer is the most frequent cancer type in women worldwide, causing about 700,000 deaths per year[1]. Most of these fatalities result from metastasis[2], a multi-step process in which cells from the tumour disseminate and colonize distant organs. Previous work has shed light on the different stages undergone by these cancer cells: invasion of the tissue surrounding the tumour, intravasation and dissemination as circulating tumour cells, extravasation, and colonization of the distant site. This process[3] involves phenotypic changes that increase the resistance of specific cells to the conditions of the “foreign” environment and result in metastasis[4]. Important in this regard is the plasticity and stemness of cancer cells, which reversibly result in epithelial, mesenchymal or stem cell-like states[5]. Secondly, the inherent heterogeneity of cancer cell populations[6],[7] at the genetic, epigenetic and microenvironmental levels predisposes some cells to the foreign environment. Thus, identifying the molecular mechanisms underlying metastatic progression is paramount to understanding this currently incurable disease and improving patient care. New technologies have been leveraged to better characterize the drivers of metastasis at various stages of the cascade, but some remain elusive due to the lack of granularity of bulk-sequencing approaches.

## Results

To better understand heterogeneity at the single-cell level, we orthotopically implanted four patient-derived xenografts (PDX)[8] and a cell line (see supplementary table) known for their lung metastatic potential into NOD-SCID-Il2rg^null^ (NSG) mice (Fig. 1a). Tumours were resected from the mammary fat pad and the animals were monitored for metastasis. Once the animals showed signs of distress (i.e., weight loss, difficulty to breath), lungs presenting metastatic lesions were collected and processed for single-cell transcriptional profiling. To exclude murine cells from the downstream analysis, human cancer cells from the tumours and matched lung metastases were purified via FACS gating GFP-positive MDA-MB-231 cells (Fig. 1b) or CD298-positive cells for PDX models (Fig. 1c). Single cells were isolated using a microfluidic device (Fluidigm C1) ahead of library preparation and sequencing. This workflow yielded a total of 1,523 single cells (Supplementary Fig. 1a) after RNA-Seq and quality control.

**Figure 1.**
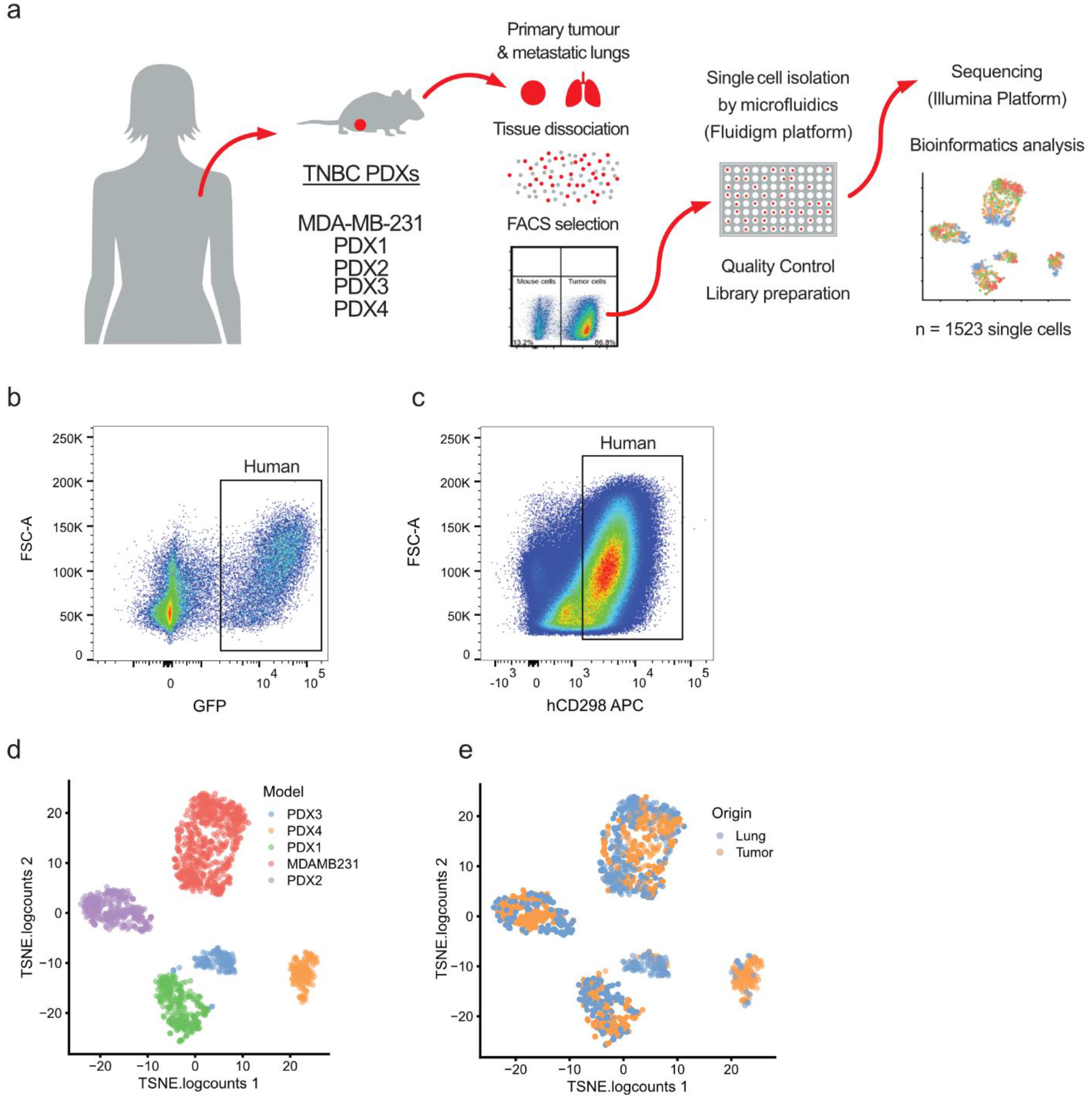
Interpatient heterogeneity is dominant over intrapatient heterogeneity. **a**, Overview of the experimental setting and models used. Human breast cancer models were implanted in the mammary fat pad of NSG mice. Tumours and lung macrometastases were harvested and mechanically and enzymatically dissociated. Human cells were purified by FACS using GFP or CD298 staining. Single cells were isolated with the Fluidigm C1 microfluidic platform and then sequenced. **b**, Representative FACS strategy for the isolation of MDA-MB-231 GFP positive. **d**, tSNE plot showing initial clustering of all the sequenced cells, according to models of origin. **e**, tSNE plot showing initial clustering of all the sequenced cells, according to the site of origin.

Initial clustering of the quality-controlled data revealed that the cells formed groups (clusters) according to the donor models (Fig. 1d). Within each cluster, cells did not clearly separate based on their origin, tumour, or metastasis (Fig. 1e). These observations highlight the importance of interpatient over intrapatient heterogeneity.

We then asked whether cells gather within each donor-specific cluster by known biological or technical features. We projected the percentage of detected genes (PDG) in each single-cell library (Fig. 2a) onto tSNE and found that the PDG influences the clustering of the cells within each given model. This variable potentially represents both biological and technical effects. Next, we assessed whether the cell cycle stage influenced the analysis of the data, as this biological variable has a broad impact on gene expression[9]. We developed a method to infer cell cycle stages in single cells (in r package gripgh) and each cell was labelled with one of the four labels (G1, G1/S, S/G2, G2/M). We found that the cell cycle stage does influence the clustering of the cells within each model (Fig. 2b).

**Figure 2.**
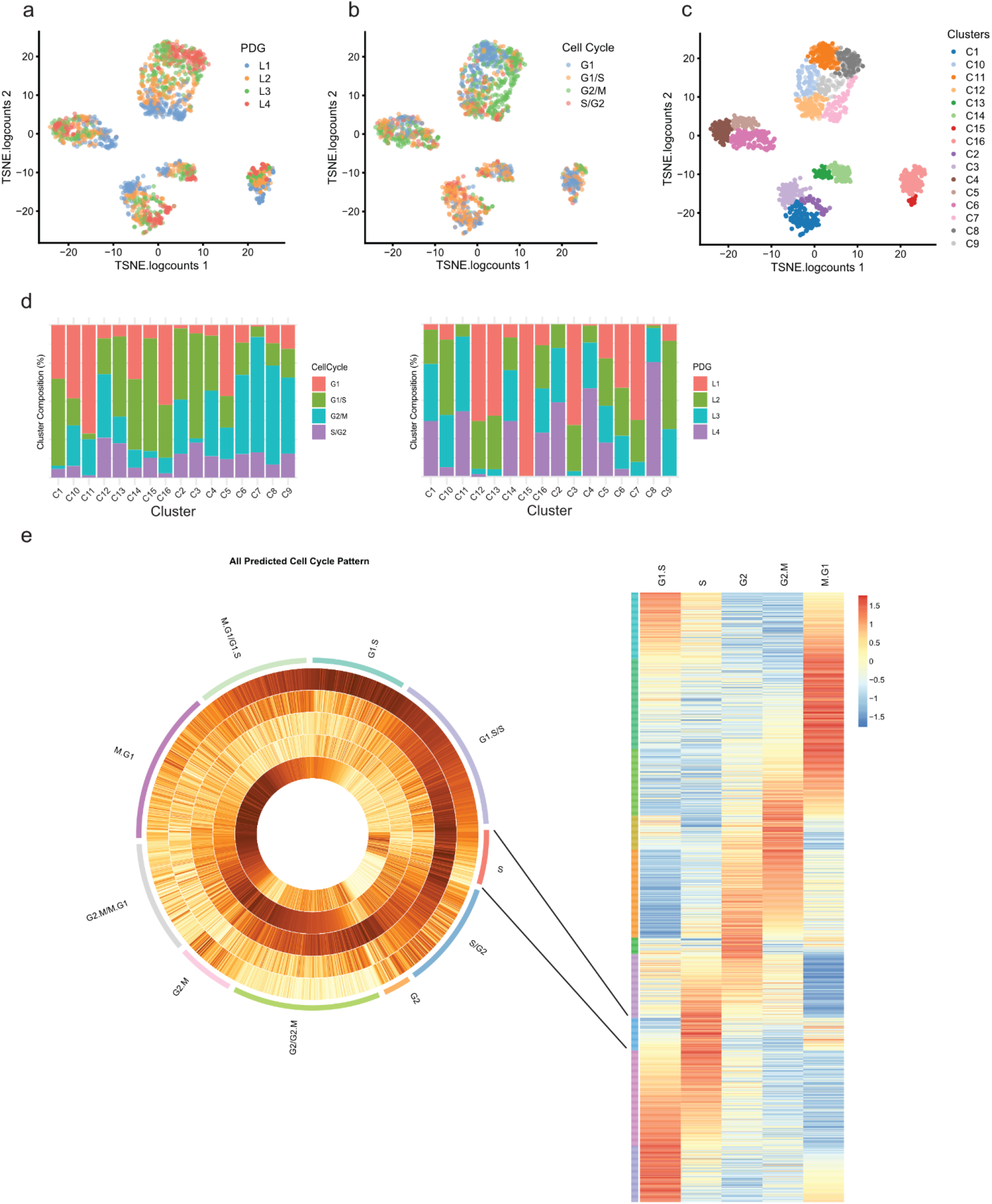
Cell cycle and percentage of detected genes delineate cell clustering. Single cells within PDX models cluster by **a**, library complexity (PDG) and **b**, cell cycle similarity. **c**, Clustering performed via t-SNE produced 16 cell clusters. **d**, Cluster composition according to cell cycle (left) and library complexity (right) **e**, An in-silico cell cycle scoring prediction to arrange single cells on a continuous cell cycle spectrum in addition to distinct cell cycle stages.

We then applied graph-based clustering to the different models and obtained 16 subclusters (Fig. 2c). These subclusters were mostly composed of cells in a similar cell cycle stage (Fig. 2d, left bar graph) and with similar PDGs (Fig. 2d, left bar graph). The influences of both the cell cycle and the PDG were also observed when the models were analysed individually (Supplementary Fig. 2a, top bar graphs for each model). Marked interpatient heterogeneity also led to the clustering of the cells according to models (Supplementary Fig. 2b). Altogether the data suggest that, in contrast to the site of origin (i.e., primary tumour or metastasis), the PDG and the cell cycle both influence the clustering of cells (Fig. 1e).

For the cell cycle stage prediction we considered that the cell cycle is not composed of discrete stages but is more a continuum of states. Gene expression is gradually modulated as a cell progresses in the cycle. Using this riche single-cell RNA sequencing data and genesets whose expression varies in different cell cycle stages, we created a circular trajectory and placed the cells in a cycle (Fig. 2e). This precise allocation along the cell cycle continuum allowed precise cell cycle staging.

To observe underlying biological processes involved in different tumour cells and to (beyond donor effect, PDG and cell cycle stage) and group biologically similar cells, we needed to remove the confounding factors of donor effect, PDG and cell cycle stage. To remove biases attributed to these factors, we used a generalized linear model (GLM)[10], correcting gene expression according to the position of the cell on the cell cycle spectrum and the complexity of the RNA-Seq library that it yielded (Fig. 3a).

**Figure 3.**
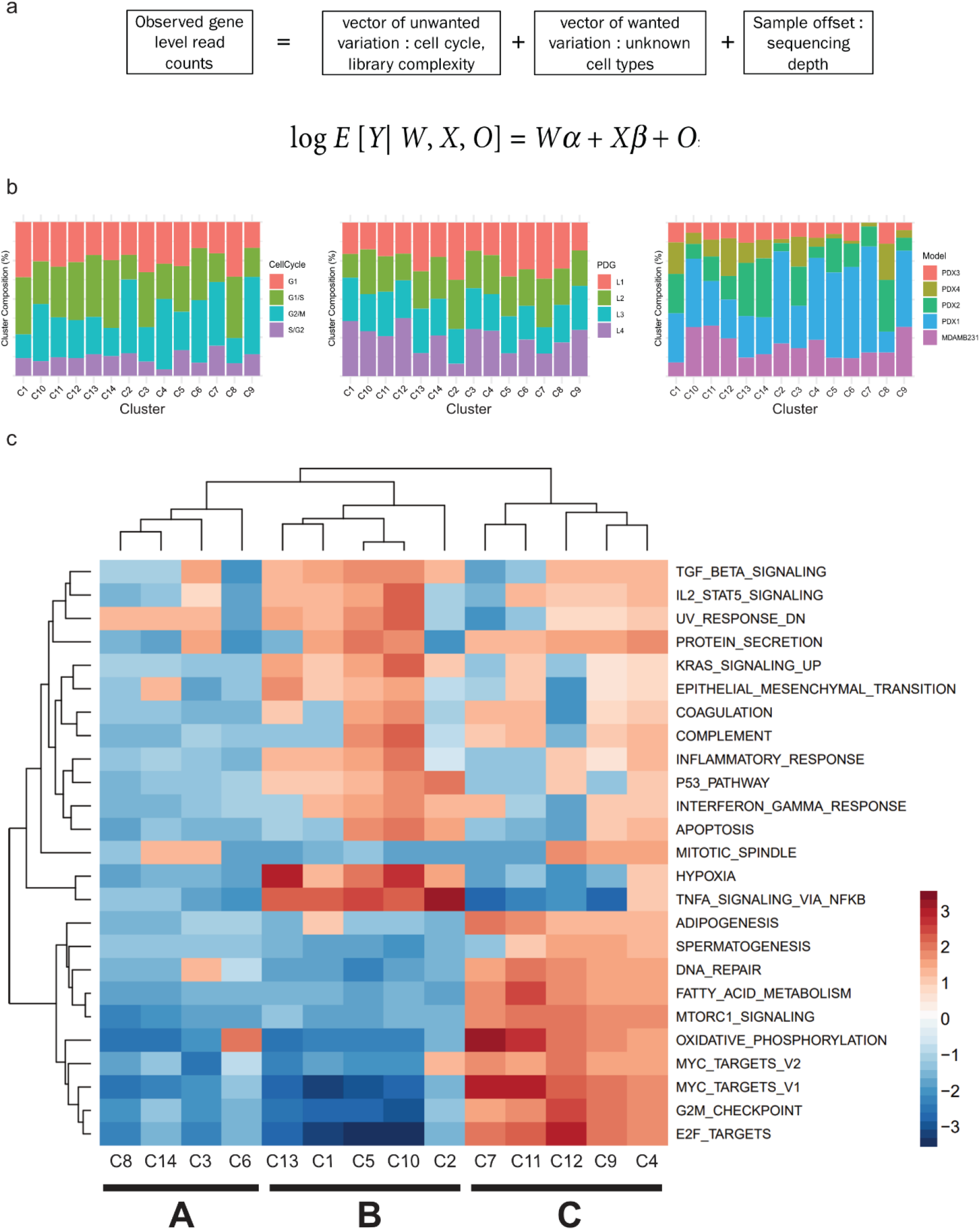
Removal of cell cycle variation and percentage of detected genes reveals three major biological clusters. **a**, Generalized linear model approach used to remove biases (cell cycle, and library complexity). **b**, Post-correction the new cell clusters have a more balanced distribution of cells from different cell cycles, library complexity (PDG), and cell source (PDX model). **c**, Gene Set Enrichment Analyses (GSEA) of individual clusters reveal common and different Hallmark genesets enriched among clusters. Top 25 of these genesets according to the absolute values of the normalized enrichment score (NES) are shown and were used to identify the superclusters.

Initially we performed gene set enrichment analysis (GSEA) with the Hallmarks gene set[11] (Supplementary Fig. 3a) on clusters defined for each model on the corrected data (Supplementary Fig. 2a, bottom bar graphs for each model, named Ax, Bx, Fx, Dx, Ex). We observed that cell clusters from different models show enrichment in a common set of biological processes. This suggests that each model contains cells in closely related biological states that could now be visualized after bias correction.

We then analysed the cells irrespective of their models or sites of origin. Using the corrected gene expression data, the cells formed 14 new clusters (Cx). The compositions of these groups were then analysed according to cell cycle stage (Fig. 3b, left), PDG (Fig. 3b, centre), and model (Fig. 3b, right). Correcting for these variables led to unbiased clustering of the cells, with clusters composed of cells from various cell cycle stages, library complexities, and models. We also plotted the composition of each cluster in terms of site of origin before and after correction (Supplementary Fig. 3b). Corrected clusters exhibited a more balanced composition, with roughly equal proportions of cells originating from the tumour and lung metastases. The data suggest that populations of cells clustering together due to their biological similarities can be found in both the tumour and the metastatic sites, without specificity to one or the other.

To further investigate the different biological states suggested by the data in Supplementary Fig. 3a, we performed GSEA on the 14 clusters formed by all the cells, regardless of their model of origin. The 14 clusters formed three “super” biological clusters (Fig. 3c). Supercluster A (C8 to C6, Fig. 3c left) is characterized by low enrichment for most of the Hallmark geneset pathways. Supercluster B (C13 to C2, Fig. 3c centre) displays the most heterogeneous regulatory landscape, with highly, moderately, and minimally enriched processes. This supercluster is defined by highly enriched Hypoxia and TNFα signalling via NFκB. The moderately enriched pathways include relevant processes such as EMT, TGFβ signalling, Interferon Gamma response, or the P53 Pathway. Finally, the least enriched genesets include MYC signalling, G2M checkpoint, and E2F targets. Interestingly these processes are most enriched in supercluster C (C7 to C4, Fig. 3c right), which also displays marked enrichment of genesets pertaining to oxidative phosphorylation, mTOR signalling, fatty acid metabolism, and DNA repair. Genesets enriched in supercluster B suggest a phenotype related to EMT, while cells in supercluster C appear to be proliferating while still partially enriched for EMT-related pathways.

We then plotted the repartition of the cells according to supercluster allocation (Supplementary Fig. 4a), noticing that cells from superclusters B and C were the most distant, with cells from supercluster A in between. We also assessed repartition according to the site of origin (Supplementary Fig. 4b) and once again found a relative balance between tumour origin and lung metastases origin of the cells forming the superclusters.

Next, we selected EMT and Proliferation markers significantly altered (FDR<0.05) in pairwise comparisons between the superclusters (Fig. 4a). Proliferation markers Ki67, MCM3, and PCNA[12] confirmed that supercluster C is the most proliferative, followed by cluster A, while supercluster B expresses these markers the least. EMT markers indicated that this process was taking place at varying levels across the different superclusters. Supercluster A displayed the least engagement in the transition according to its low expression of several EMT markers (ZEB1, SOX9, SNAI1, FN1, TGFBR1). Superclusters B and C showed increased expression of these markers but at varying levels. Such heterogeneity suggests that these superclusters may undergo EMT but could be at different stages of the process. Such partial EMT has previously been described[13], [14] and may reflect the balance between proliferative potential and migratory capability, both properties being typical of different stages of the metastatic cascade.

**Figure 4.**
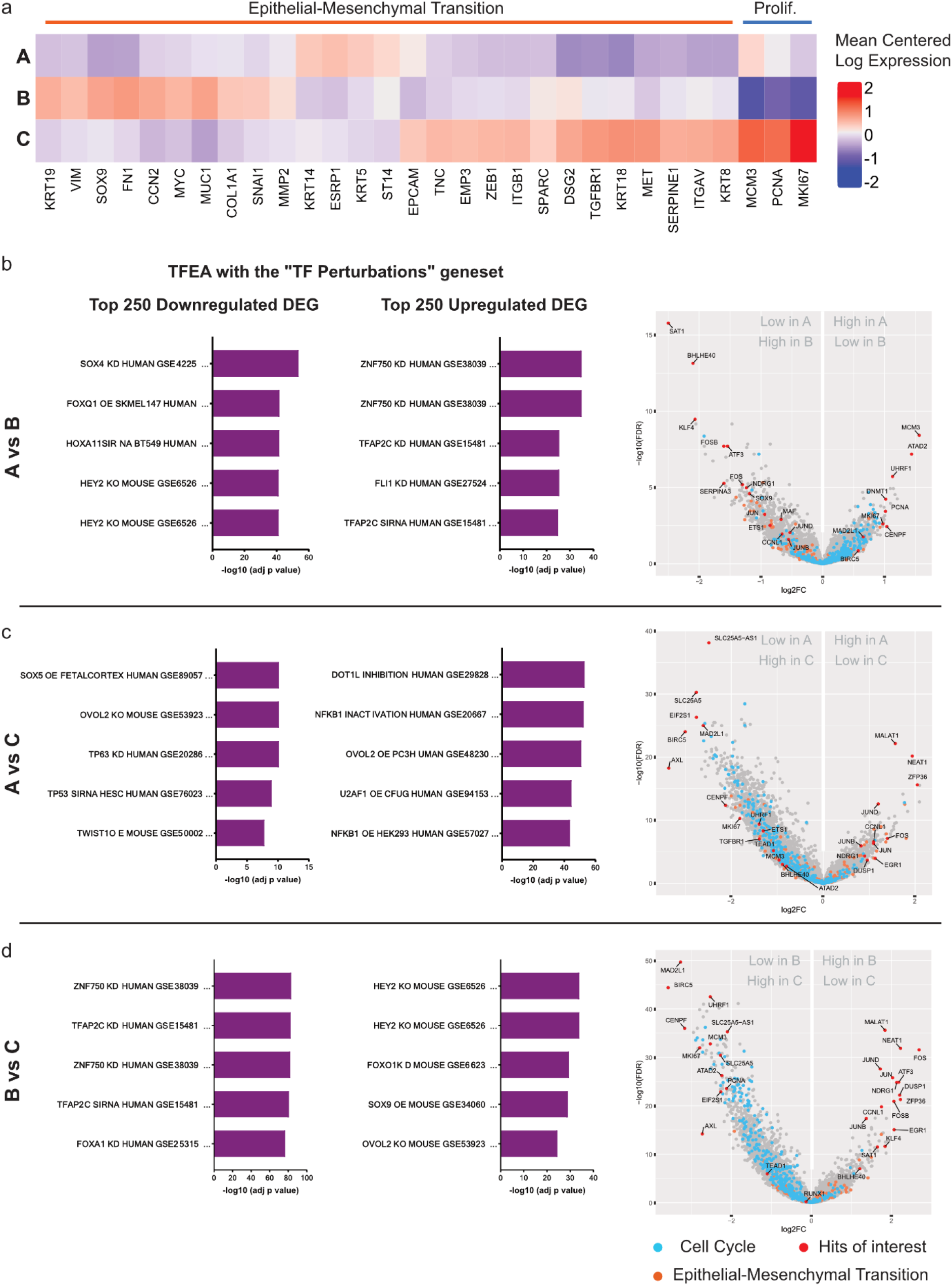
Comparison of major biological superclusters reveals partial EMT state regulators. **a**, EMT and Proliferation markers significantly altered (FDR<0.05) in supercluster pairwise comparisons. **b, c** and **d**, Right: Transcription Factors Enrichment Analysis of the top 250 up- and downregulated differentially expressed genes of each supercluster. Left: Volcano plots highlighting the cell cycle gene set (blue, Bioplanet 2019), the EMT gene set (orange, mSigDB) and hits of interest (red) in corresponding supercluster pairwise comparisons.

To investigate partial EMT states of the superclusters, we selected the top 250 up- and downregulated genes of each cluster and performed GSEA as well as Transcription Factor Enrichment Analysis (TFEA) with the EnrichR[15] platform (Fig. 4a, b, c bar plots).

The “TF Perturbations followed by expression” geneset, which was generated by the curation of experiments altering TFs before measuring gene expression, revealed that different TFs govern the top differentially expressed genes (DEGs) across the superclusters. These uncommon TFs paint a complex picture of the EMT states existing in superclusters A, B, and C. HEY2, OVOL2, TFAP2C, and TP63 are TFs described as regulators of partial EMT states in mouse models[14]. ZNF750 was recently described as an EMT repressor in breast cancer[16], an activity shared by OVOL2[17], TP63[14], and FOXO1[18]. HEY2, TFAP2C[19] [20], SOX4, and SOX9[21] are known to promote EMT. U2AF1 is a splicing factor fine-tuning translation with reported effects in development and EMT.

These TFs control the EMT and proliferation state of the superclusters shown in Fig. 4a. Supercluster A, shown to be mildly proliferative, is controlled by TFs evocating differentiated slowly proliferating cells. E2F4 is known to be abundant in differentiated cells and to repress proliferative genes[22]. TP63 and TFAP2C have been described as controlling early hybrid EMT states, with cells close to an epithelial state and more prone to proliferate than mesenchymal cells. Additionally, KDM5B has been reported to characterize a slow-cycling cell subpopulation in melanoma[23], which fits the traits of supercluster A. Supercluster B, the least proliferative, is also the one with the strongest expression of canonical EMT markers/TFs such as FN1, SNA1, MYC, and SOX9. When compared to superclusters A and C, DEGs from supercluster B appear to be under the control of HEY2, a member of the basic Helix-Loop-Helix (bHLH) transcription factor family. bHLH have been described as late hybrid EMT regulators[14] responsible for mesenchymal phenotypes that favour migration and have little proliferation potential. Supercluster C, the most proliferative, is controlled by ZNF750, TFAP2C, and OVOL2. These TFs have been described as either EMT repressors or regulators of early hybrid EMT stages, corresponding to epithelial-like phenotypes permitting proliferation. It should however be noted that the EMT markers are high in this supercluster, indicating that the process is ongoing yet likely oriented towards the proliferation of cells rather than their migration.

Next, we analysed individual hits from the pairwise supercluster comparisons (Fig. 4a,b,c, volcano plots). Several members of the Activator Protein 1 (AP1) family of transcription factors (JUN, FOS, ATF3) were consistently altered in the different superclusters. This is of importance as AP1 has been reported as one of the “core” TFs binding enhancers regulating epithelial and mesenchymal states[14], [24]. This core is subsequently modulated by other TFs such as those described in the previous paragraph. Another element strongly differing between superclusters was BHLHE40, a member of the bHLH TF family, which was found strongly downregulated in supercluster A. BHLHE40 has been reported to induce EMT as well as tumour growth and lung metastases via HBEGF exosomal release[24]. Different types of RNA-coding genes are modulated during EMT, some of which (NEAT1, MALAT1) figured in the top DEGs across the superclusters. NEAT1 was found upregulated in superclusters A and B compared to cluster C. NEAT1 has been described as promoting chemoresistance and cancer stemness[25]–[27]. More importantly, it was found to enhance glycolysis as a scaffold of key glycolytic enzymes. Its downregulation in supercluster C, which is enriched for fatty acid metabolism and the oxidative phosphorylation process, is notable. This observation may reflect a metabolic shift away from glycolysis and towards oxidative phosphorylation that was shown to exacerbate breast cancer lung micrometastases. Like NEAT1, MALAT1 is upregulated in superclusters A and B compared to C. Its effects are manifold and some controversy exists about its activities in different cancer types and settings[28], [29]. It is also interesting to note that MALAT1 has been described to interact with TEAD, which is part of the core TFs controlling the EMT process.

Finally, to relate each of the biological states to patient outcome, we selected the upregulated transcripts characteristic of each supercluster by overlapping the different contrasts (see Supplementary Table 2). The transcripts defining supercluster A correlate with better relapse-free survival (RFS) in patients suffering from basal-like breast cancer (Supplementary Fig. 4c). On the contrary, upregulated transcripts found in superclusters B and C (Supplementary Fig. 4d and 4e respectively) correlated with worse RFS. These results suggest that these cell states may exhibit different levels of aggressiveness driven by different sets of transcripts.

## Discussion

The cellular plasticity arising from the transitions between epithelial and mesenchymal states has been recognized as instrumental to metastatic progression[5]. The characterization of cancer cells at different stages of the metastatic cascade in terms of canonical EMT markers and TFs has recently given rise to the concept of partial EMT[10, 27, 28]. The balanced epithelial and mesenchymal traits in partial EMT states results in positive reactions to the conditions imposed by the metastatic process[31] such as dissemination as CTCs or colonization of a foreign microenvironment. Characterization of these partial EMT states is however delicate, model- and context-dependent, and a task for which the use of canonical EMT markers and TFs is not sufficient.

Using scRNA-Seq on tumours and lung macrometastases of PDX models of TNBC, we observed marked heterogeneity at the intra- and interpatient levels. After correcting for the two confounding factors cell cycle and PDG, we identified three cell states (superclusters) present at both sites. These superclusters differ from each other in EMT and proliferation markers. Analysis of the top DEGs of each supercluster revealed that the variation between them is controlled by a set of TFs (i.e., TFAP2C, ZNF750, OVOL2, TP63, HEY2, bHLHs), which have been recently shown to finely modulate partial EMT states in other models[32]. These TFs act by modulating core TFs such as AP-1 (composed of its JUN and FOS subunits), which we also found deeply altered across the superclusters. Our findings highlight the presence of partial EMT states controlled by the aforementioned TFs in breast cancer PDX models, both in the tumour and lung macrometastases. While our data have not yet allowed identification of factors driving lung macrometastases specifically, they still shed light on breast cancer tumour and metastases biology at this specific stage. Additionally, our results highlight the importance for breast cancer of several elements such as NEAT1 or MALAT1. These lncRNAs have been recently implicated in breast cancer initiation, growth, metastasis, and chemoresistance[23, 30–33]. Their downregulation in the most proliferative supercluster questions their roles in these cells and whether choosing them as therapeutic targets could affect the different cell states identified here. Such dynamics may ultimately affect how tumours and metastases respond to treatment. Our results highlight the marked heterogeneity of breast cancer cells. They call for further studies at the single-cell level to better characterize the different partial EMT states at different stages of the metastatic cascade. Future studies, especially those including the stromal and immune compartments, will further our knowledge of drivers of metastasis and how to tackle them.

## Supplementary Information

## Acknowledgements

We thank the members of the Bentires-Alj team who participated in discussions about this project. We are also grateful to the core facilities of the FMI and DBM for supporting this work. We also thank Tim Roloff for his insights into single-cell RNA-seq. The Bentires-Alj laboratory research is supported by the Swiss Initiative for Systems Biology (SystemsX), the Swiss National Science Foundation, the European Research Council (ERC advanced grant 694033 STEM-BCPC), the Krebsliga Beider Basel, the Swiss Cancer League, the Swiss Personalized Health Network, and the Department of Surgery of the University Hospital Basel.

## Author contributions

B.H analysed the data, interpreted the results, and wrote the manuscript. M.M.S.O conceived the study, designed and performed the experiments, and analysed the data. S.M performed histopathological analysis of the PDX models. A.S performed computational analysis, designed the cell cycle/PDG correction method, analysed the data and interpreted the results. M.K performed data analysis and result interpretation. C.B and K.E, helped with the experimental design and use of the single-cell platform. H.K performed fluorescence-activated cell sorting experiments, analysis, and interpretation of results. M.S advised on experiment design and data analysis. S.S, M.V, W.P.W provided patient material. M.B-A conceived the study, designed the experiments, and interpreted the results. All authors read and provided feedback on the manuscript.

## Author information

B.H, M.M.S.O, A.S, M.K declare no competing interests. M.M.S.O and A.S are employees of Roche. M.B.-A. owns equities in Novartis.

## Materials and correspondence

Correspondence and requests for materials should be addressed to Mohamed Bentires-Alj.

This manuscript contains 4 Figures, 4 Extended Data Figures, 2 Supplementary Tables.

## Methods

### *In vivo* experiments

All *in vivo* experiments were performed in accordance with the Swiss animal welfare ordinance and were approved by the cantonal veterinary office, Basel Stadt. Female NSG and BALB/c mice were maintained in the Friedrich Miescher Institute for Biomedical Research and the Department of Biomedicine animal facilities in accordance with Swiss guidelines on animal experimentation. Mice were maintained in a sterile environment with light, humidity, and temperature control (light-dark cycle with light from 7:00 to 17:00, with a gradual change from light to dark, temperature 21–25 °C, and humidity 45–65%). Before each experiment, mice were allowed to acclimatize for a minimum of seven days. MDA-MB 231 cells (10,000 cells) were resuspended in 40 μl Matrigel:PBS (1:1) and injected into the pre-cleared mammary fat pads of 4- to 8-week-old female NSG mice. PDX models were transplanted into the pre-cleared 4th mammary fat pads of NSG mice. Tumours were resected when the largest diameter reached 10 mm and mice were monitored regularly for signs of metastatic outgrowth and distress. In none of the experiments did tumour volumes exceed approved limits. All orthotropic experimental procedures (tumour resection and tumour cell implantation) were undertaken on anesthetized mice by a single investigator, according to protocols approved by the cantonal veterinary office Basel Stadt. Experimental metastasis assays were performed by injecting 100,000 cells into tail veins. After intravenous injection of MDA-MB 231 cells, we performed *in vivo* bioluminescence imaging to confirm injection and to monitor metastatic outgrowth. Bioluminescence imagining was performed using an IVIS Lumina XR (Caliper LifeSciences) upon injection of luciferin (Biosynth; L8220).

### Cell lines and PDX models

The cell lines MDA-MB 231 andHEK293T were purchased from ATCC and cultured according to ATCC protocols. Cell line identity was confirmed and routinely tested using short tandem repeat sequencing; all cell lines were routinely tested for mycoplasma contamination. MDA-MB 231 were propagated in monolayer cultures in DMEM supplemented with 10% FCS. All experiments were performed with 70–90% confluent cells. The PDXs used in this study have previously been described[37]. PDX models were passaged in NSG animals via tumour piece implantation in the 4th mammary fat pad. Immunohistochemistry staining was performed to ensure the ER/PR/HER2 status.

### Lentiviral vectors, lentivirus, and infection

Lentiviral batches were produced using PEI transfection on HEK293T cells as previously described[38]. The titre of each lentiviral batch was determined in MDA-MB 231 cells. Cells were infected for 8 h in the presence of polybrene (8 μg/ml). Selection with 2 μg/ml puromycin (Sigma) was applied 48 h after infection.

### Sample preparation

Tumours and matched lung metastases were dissociated into single cells using mechanical disruption followed by enzymatic digestion by a collagenase-hyaluronidase solution (StemCell Technologies; 07912) for 1 h at 37°C without mechanical agitation. The resulting material was filtered twice on 40-μm cell strainers and depleted of erythrocytes using a red blood cell lysis buffer (Sigma, R7757).

### Fluorescence-activated cell sorting

MDA-MB-231 expressing a GFP-Luciferase construct were selected based on GFP expression. PDX models cells selection relied on the CD298 human-specific marker. Sorts were performed on a BD FACS BD Aria III (70μm nozzle). DAPI staining was used to gate out dead cells. Single cells and doublets were respectively and excluded based on their forward and side scatter profiles and pulse width. The APC anti-human CD298 antibody (Biolegend, 341706) was used.

### scRNA-Seq

Human cells were processed for single cell isolation and library preparation using the Fluidigm C1 platform. Single-cell capture was performed by microfluidics on medium and large Fluidigm C1 integrated fluidics chip (IFC) (Fluidigm; PN100-5760 and PN100-5761). Visual quality control by microscopy allowed assessment of capture efficiency. cDNA was generated from the captured cells as per the manufacturer’s protocol using SMART-Seq Ultra Low RNA (Takara Bio; 634833) before being processed for Illumina sequencing via the Nextera XT DNA Library Preparation kit (Illumina; FC-131-1096). Sequencing was performed on an Illumina platform.

### Computational analysis

#### RNA sequencing data analyses

Reads were aligned to the human genome (UCSC version hg38AnalysisSet) with STAR. The output was sorted and indexed with samtools. Strand-specific coverage tracks per sample were generated by tiling the genome in 20-bp windows and counting 5’end of reads per window using the function bamCount from the bioconductor package bamsignals. These window counts were exported in bigWig format using the bioconductor package rtracklayer. The rsubread::featureCounts function was used to count the number of reads (5’ends) overlapping with the exons of each gene, assuming an exon union model (gene annotation: ensembldb_Homo_sapiens_GRCh38_ensembl_96.sqlite).

#### Removal of potential doublets

Following observation under the microscope, wells containing more than 1 cell or debris were marked and removed from further analyses. To further remove potential human and mouse multiplets, we used fastq_screen to count reads mapping uniquely to human and mouse genomes (human-to-mouse ratio). Libraries with a human-to-mouse ratio >=5 were considered human cells and retained for further analyses. Libraries with <= 100 k reads mapping to human transcriptome were also filtered out. Libraries expressing less than 2,346 genes and more than 9,884 genes were also filtered out, retaining the cells that expressed 10% - 40% of the total number of unique genes observed in all models (23,459 genes). This step removed libraries with low complexity and a few outlier libraries where very high numbers of genes were observed.

#### Inference and correction of cell cycle signal

The cell cycles of individual cells were inferred using sets of cell cycle-regulated genes known to peak in transcription at given cell cycle stages obtained from (PMID: 12058064, PMID:11416145). For each cell, five normalized cell-cycle stage scores (G1.S, G2, G2.M, M.G1, S) were calculated and the cell assigned to that most closely resembling the expected profile based on correlation. Optionally, labels were further refined by iteratively estimating new cell cycle score profiles based on estimated cell labels, and re-assigning cells to the profile with the highest correlation. The function call predictCellCycle(org = “human.Whitfield”, cor_thr = 0.2, refine_iter = 200) is available as part of R package griph (https://github.com/ppapasaikas/griph). The cell cycle and library complexity (percentage of detected genes) were modeled as covariates and regressed out of the log-normalized counts using glm.fit (R package glm).

#### Dimensionality reduction, clustering, differential gene expression

Each library was normalized to 100-K reads and log transformed adding pseudocount of 1. PCA, tSNE, and UMAP projections (on log-normalized data or residuals) were computed with R package scater using default parameters. Nearest-neighbour graphs were computed using function buildSNNGraph (R package scran) with tSNE and UMAP as inputs separately. Function cluster_louvain (R package igraph) was used for graph-based clustering. Differential gene expression between single-cell clusters was performed using pairwise t-test implemented in FindMarkers function from R package scran. The values pval.type=“some”, min.prop=0.2 were used as arguments in FindMarkers, indicating that the genes considered as marker genes are those differentially expressed in at least 20% of the pairwise comparisons, i.e. combined p-value from the pairwise comparisons was calculated by taking the minimal value of the top 20% Holm-corrected p-values for each gene. Genes were ranked by the respective combined logFC (summary.logFC) and gene set enrichment was performed with bioconductor packages fgsea and msigdb (collections H, C2, C5).

### RFS Analysis

Kaplan-Meier plots were generated from the KMplotter database using the mRNA Gene Chip dataset. Upregulated transcripts specific to a supercluster were isolated (with the Venny online tool) from the different contrasts generated from the analysis of the Top 250 DEGs used for the rest of the analysis. The generated lists then served as input to the KMplotter tool via the Mean Expression for Multiple Genes function. RFS analysis was performed on the patients with basal-like breast cancer (PAM50 classifier, *n*=442) and with the Autoselect Best Cutoff parameter.

## Data availability

The sequencing data have been deposited in the Gene Expression Omnibus (GEO) and are accessible under the accession code GSE202695. Processed transcriptomic data is available upon request to the corresponding author.

