## Supplementary Figures for "Single-cell analysis reveals inter- and intratumour heterogeneity in metastatic breast cancer"

### Extended Figure 1

a

| Model | Tumor | Lung Metastases |
|---|---|---|
| **PDX4** | 151 | 17 |
| **PDX3** | 23 | 115 |
| **PDX2** | 177 | 114 |
| **PDX1** | 298 | 314 |
| **MDA-MB-231** | 221 | 93 |
| **Total** | 870 | 653 |

#### Extended Figure 2

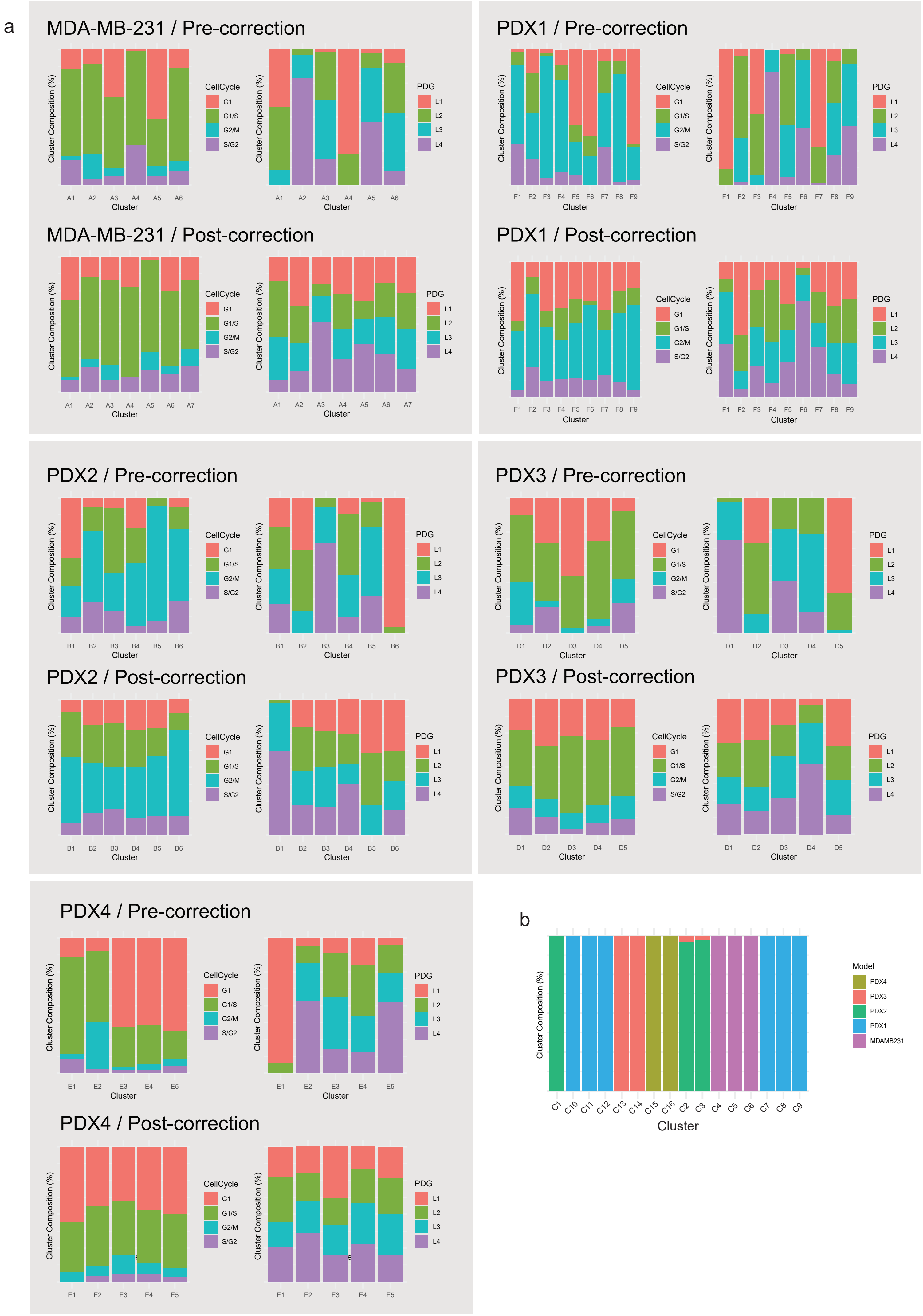

#### Extended Figure 3

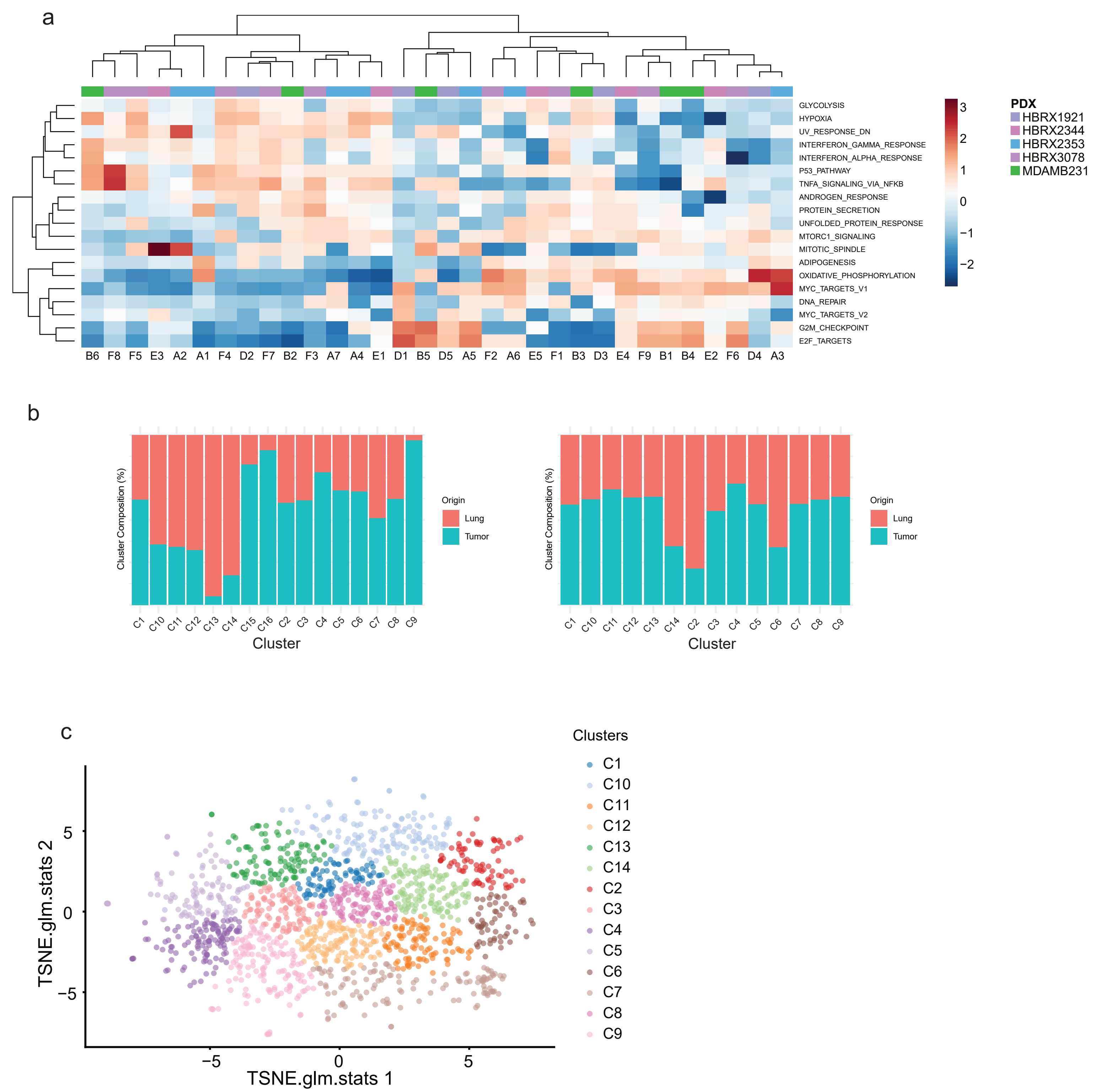

**Extended Figure 4**

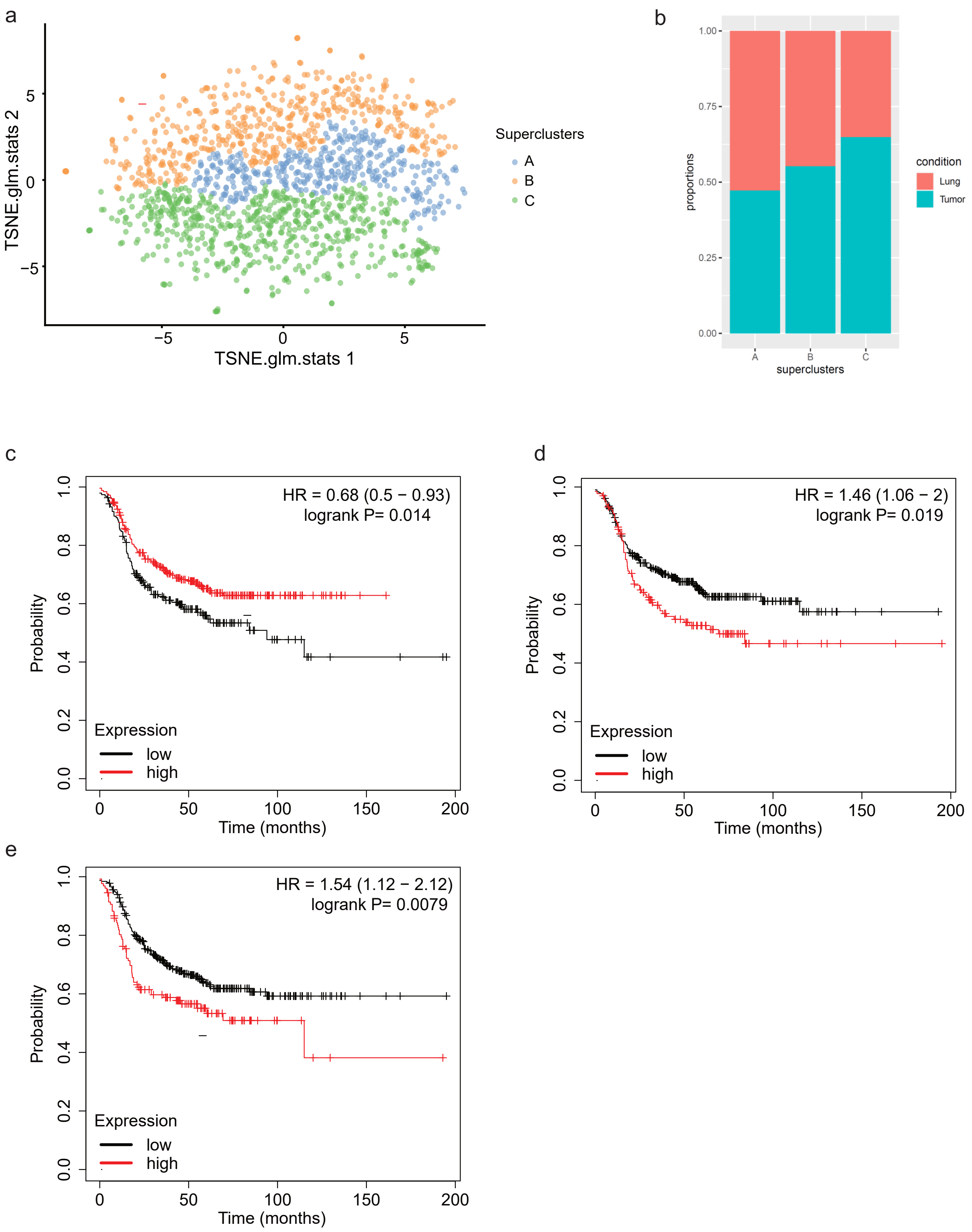

**Extended Figure 1**

**a, Table summarizing the numbers of cells sequenced and processed for analysis for each model / site of origin after quality control**

**Extended Figure 2**

**a, Clustering and distribution of cells from different cell cycle stages and library complexity (PDG) before correction (top row) and after correction (lower row) for each model.**

**b, Clustering performed over tSNE produces 16 cell clusters before bias correction**

**Extended Figure 3**

**a, Cell clusters identified separately from each individual model, show overlapping enrichment in Hallmark genets**

**b, Corrected clusters generated from all the cells irrespectively of the model of origin, showing the cluster composition according to site of origin**

**c, tSNE plot showing clustering of the corrected clusters**

**Extended Figure 4**

**a, tSNE plot showing the cell repartition across the 3 superclusters defined via GSEA**

**b, Barplot  showing the supercluster composition according to site of origin**

**c, Kaplan-Meyer plot showing the RFS in patients with basal-like BC (n=442) according to expression of the upregulated transcripts defining supercluster A**

**d, Kaplan-Meyer plot showing the RFS in patients with basal-like BC (n=442) according to expression of the upregulated transcripts defining supercluster B**

**e, Kaplan-Meyer plot showing the RFS in patients with basal-like BC (n=442) according to expression of the upregulated transcripts defining supercluster C**
